# The Two Amino Acids Proximate to the Renin Cleavage Site of Human Angiotensinogen Do Not Affect Angiotensin II-mediated Functions in Mice

**DOI:** 10.1101/2019.12.18.875914

**Authors:** Chia-Hua Wu, Congqing Wu, Deborah A. Howatt, Jessica J. Moorleghen, Lisa A. Cassis, Alan Daugherty, Hong S. Lu

## Abstract

**Objective:** Renin cleavage of angiotensinogen (AGT) has species specificity. Since the residues at positions 11 and 12 of AGT are different between human and mouse AGT, we determined whether these two residues in AGT affect renin cleavage and angiotensin II-mediated functions using an adeno-associated viral (AAV) approach for manipulating AGT in vivo.

**Approach and Results:** Hepatocyte-specific AGT deficient (hepAGT-/-) mice in an LDL receptor -/- background were infected with AAVs containing a null insert, human AGT, or mouse AGT expressing the same residues of the human protein at positions 11 and 12 [mouse AGT (L11V;Y12I)]. Expression of human AGT in hepAGT-/- mice led to high plasma human AGT concentrations without changes in plasma mouse endogenous AGT, plasma renin concentrations, blood pressure, or atherosclerosis. This is consistent with human AGT not being cleaved by mouse renin. To determine whether the residues at positions 11 and 12 in human AGT lead to the inability of mouse renin to cleave human AGT, hepAGT-/- mice were injected with AAV encoding mouse AGT (L11V;Y12I). Expression of mouse AGT (L11V;Y12I) resulted in increased plasma mouse AGT concentrations, reduced renin concentrations, and increased renal AngII concentrations that were comparable to their concentrations in hepAGT+/+ mice. This mouse AGT variant increased blood pressure and atherosclerosis in hepAGT-/- mice to the magnitude of hepAGT+/+ mice.

**Conclusion:** Replacement of L11 and Y12 to V11 and I12, respectively, in mouse AGT does not affect renin cleavage and AngII-mediated functions in mice.

**HIGHLIGHTS:**

- Human AGT is not cleaved by mouse renin and does not change AngII-mediated functions in hepatocyte-specific AGT -/- mice.
- Replacement of the N-terminal amino acids at 11 and 12 positions from mouse to human AGT does not affect renin cleavage and AngII-mediated functions in hepatocyte-specific AGT -/- mice.

## INTRODUCTION

Renin cleavage of angiotensinogen (AGT) is the rate-limiting step in producing bioactive angiotensin (Ang) peptides.^1,2^ The product of AGT and renin interaction is AngI, the first 10 amino acids of the amino terminus of secreted AGT. AngI is highly conserved from fly, rodents, to human.^2^ One study reported that a conserved disulfide bond between Cys18 and Cys138 in human AGT (Cys18 and Cys137 in mouse AGT) regulated cleavage of human AGT by human renin in vitro.^3^ However, our in vivo study demonstrated that this conserved disulfide bond in AGT did not affect AngII production and AngII-mediated functions,^4^ implicating potentially different results from in vitro versus in vivo conditions.

Renin cleavage of AGT displays species specificity.^5-7^ For example, human AGT is not cleaved by mouse renin, and vice versa as demonstrated in transgenic mouse models.^6,7^ Similarly, human AGT is not cleaved by canine renin, but replacement of the residues at positions 11 and 12 of human AGT to the two residues in canine AGT enhanced its cleavage by canine renin in an in vitro study,^8^ implicating that these two residues adjacent to the renin cleavage site are important for regulating AGT cleavage by renin. However, it has not been determined whether the two amino acids adjacent to the AngI sequence in AGT affect renin cleavage and AngII-mediated functions in vivo.

In this study, we used an adeno-associated viral vector (AAV)-driven expression approach to either populate the full length of human AGT or mouse AGT that its two residues at positions 11 and 12 were replaced with the two residues in the human protein. This approach provided a rapid and efficient manipulation of AGT to study amino acid determinants of its interaction with renin in vivo and the consequent AngII-mediated physiological and pathophysiological functions.

## MATERIALS AND METHODS

Detailed Materials and Methods are available in the online-only Data Supplement. The data that support the findings reported in this manuscript are available from the corresponding authors upon reasonable request.

### Animals

Development of hepatocyte-specific AGT deficient mice has been reported previously.^4,9,10^ AGT floxed (termed “Agt f/f”) x albumin-Cre^-/-^ (hepAGT +/+) and Agt f/f x albumin-Cre^+/-^ (hepAGT -/-) littermates were used for experiments described in this manuscript. Genotypes were determined prior to weaning and validated after termination by Cre PCR, and confirmed by measuring plasma AGT concentrations during the study and after termination. Both male and female mice in an LDL receptor -/- background were studied in accord with the recent ATVB Council statement.^11^ Systolic blood pressure was measured on conscious mice using a non-invasive tail-cuff system (Coda 8, Kent Scientific Corporation) following our standard protocol.^12^ Atherosclerosis was quantified following the recommendations of the AHA statement.^13^ All animal experiments reported in this manuscript were performed with the approval of the University of Kentucky Institutional Animal Care and Use Committee (IACUC protocol number 2006-0009 or 2018-2968).

### Adeno-associated viral vectors (AAVs) encoding human or mouse AGT

AAV vectors (serotype 2/8) driven by a hepatocyte-specific thyroxine-binding globulin (TBG) promoter were produced by the Vector Core in the Gene Therapy Program at the University of Pennsylvania. Three AAV vectors were made to encode: (1) a null insertion (null AAV; used as control), (2) the human AGT (human AGT.AAV), and (3) the mouse AGT with L11V;Y12I substitutions (L11V;Y12I.AAV). The L11V;Y12I represented that the N-terminal leucine at position 11 and tyrosine at position 12 in mouse AGT were replaced by valine and isoleucine, respectively, to mimic the two amino acids at the same positions of human AGT (Figure 1A). Experimental design was summarized in Figure 1B.

**Figure 1.**
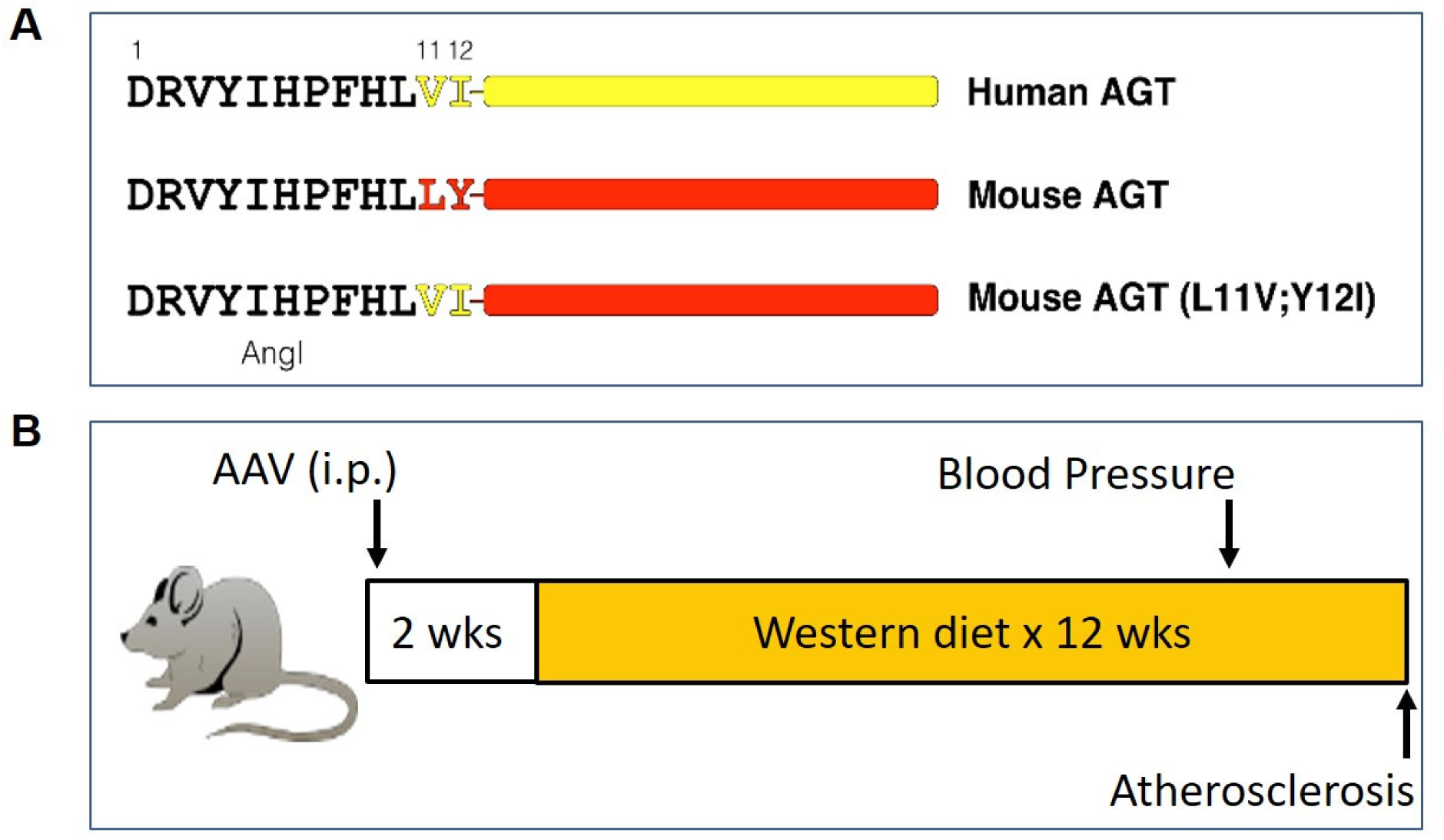
**(A)** N-terminal amino acids 1-10 (Angl) are the same between human and mouse AGT. Amino acids at positions 11 and 12 are different between human and mouse AGT. Mouse (L11V;Y12I) represent the N-terminal leucine at position 11 and tyrosine at position 12 in mouse AGT replaced by valine and isoleucine, respectively, to mimic the two amino acids at the same positions of human AGT. **(B)** Schematic summary of the experimental design: Mice were injected intraperitoneally with AAVs encoding a null insert, human AGT, or mouse AGT (L11V;Y12I). Two weeks after AAV injections, mice were fed a Western diet for 12 weeks. Blood pressure was measured on week 10 during the Western diet feeding, and atherosclerosis was measured after termination.

### Statistical analysis

Data are represented as means ± standard error of means (SEM). SigmaPlot version 14.0 (SYSTAT Software Inc.) was used for statistical analyses. To compare multiple-group data, one-way ANOVA followed by Holm-Sidak method was used for normally distributed variables that passed equal variance test. Kruskal-Wallis one-way ANOVA on Ranks followed by Dunn’s method was used for data that did not pass either normality or equal variance test. P < 0.05 was considered statistically significant.

## RESULTS

### Population of human AGT in hepAGT -/- mice did not affect systolic blood pressure and atherosclerosis

In both hepAGT +/+ and -/- male mice infected with null AAV, human AGT was not detected in mouse plasma by ELISA. In hepAGT -/- mice infected with AAV encoding human AGT, plasma human AGT concentrations were comparable to plasma AGT concentrations of humans^14^ (Figure 2A), but did not affect plasma concentrations of endogenous mouse AGT (Figure 2B), demonstrating induction of human AGT did not affect endogenous mouse AGT concentrations in hepAGT -/- mice. Plasma renin concentrations were comparable between hepAGT -/- mice infected with null AAV and human AGT.AAV, both of which were higher than those in hepAGT +/+ mice injected with null AAV (Figure 2C). Plasma total cholesterol concentrations were above 1,000 mg/dl after 12 weeks of Western diet feeding with no differences among the 3 groups (Figure 2D). Since AngII, the major effector peptide of the renin-angiotensin system, regulates blood pressure and contributes to development of atherosclerosis, we measured these two parameters. Expression of human AGT in hepAGT -/- mice did not change systolic blood pressure and atherosclerotic lesion size, when compared to hepAGT-/- mice infected with null AAV (Figure 2E and F). However, systolic blood pressure and atherosclerotic lesion size were much higher in hepAGT +/+ mice than in hepAGT -/- mice injected with either null AAV or human AGT AAV. These findings were also confirmed in female mice (Figure I in the online-only Data Supplement), demonstrating that human AGT does not affect AngII-mediated functions in mice irrespective of sex.

**Figure 2.**
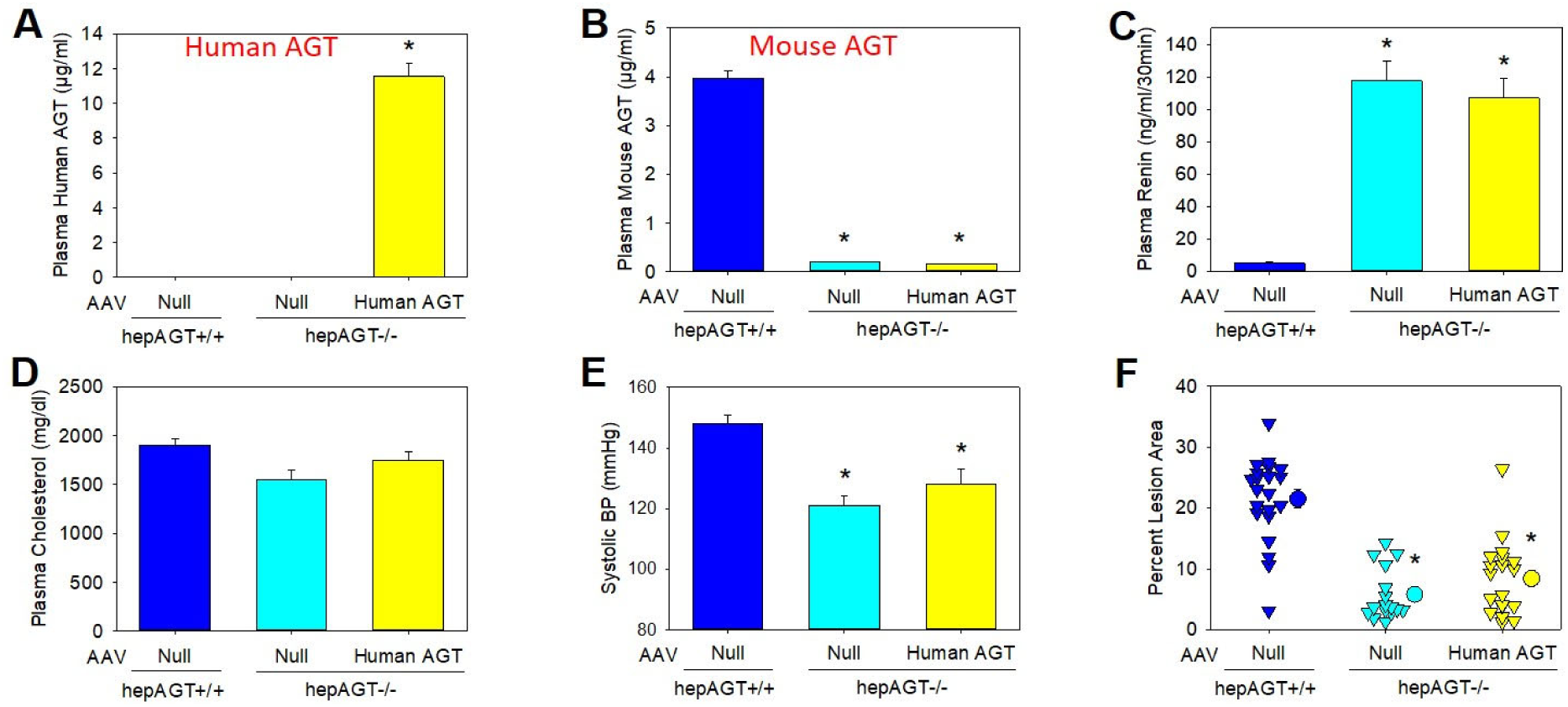
Human AGT did not affect Angll-mediated functions in male hepAGT-/- mice. **(A)** Plasma human AGT concentrations were measured by an ELISA kit that was specific for human AGT. *P<0.05 versus the other 2 groups by Kruskal–Wallis one way ANOVA on Ranks with Dunn’s method. **(B)** Plasma mouse AGT concentrations were measured by an ELISA kit that was specifically targeting mouse AGT. *P<0.05 versus hepAGT +/+ mice injected with null AAV by Kruskal–Wallis one way ANOVA on Ranks with Dunn’s method. **(C)** Plasma renin concentrations were measured by radioimmunoassay. *P<0.05 versus hepAGT +/+ mice injected with null AAV by Kruskal–Wallis one way ANOVA on Ranks with Dunn’s method. **(D)** Plasma total cholesterol concentrations were measured by an enzymatic assay kit and analyzed with one way ANOVA. P>0.05. **(E)** Systolic blood pressure was measured using a tail-cuff system 3 weeks after AAV injection. *P<0.05 versus hepAGT +/+ mice injected with null AAV by one way ANOVA with Holm-Sidak method. **(F)** Atherosclerotic lesion area was quantified by an *en face* method. *P<0.05 versus hepAGT +/+ mice injected with null AAV by one way ANOVA with Holm-Sidak method. N = 16 - 22/group.

### Replacement of amino acids 11 and 12 in mouse AGT restored AngII production and AngII-mediated functions in hepAGT -/- mice

To determine whether N-terminal residues at positions 11 and 12 of AGT affect renin cleavage and AngII-mediated functions, AAV encoding mouse AGT with the L11V and Y12I substitutions were injected into male hepAGT-/- mice. Administration of mouse AGT(L11V;Y12I).AAV led to increased plasma mouse AGT concentrations to a comparable level as in hepAGT +/+ mice injected with null AAV (Figure 3A). Plasma renin concentrations were not different between hepAGT +/+ mice and hepAGT -/- mice populated with mouse AGT(L11V;Y12I), but lower than hepAGT -/- mice infected with null AAV (Figure 3B). As reported previously,^10^ plasma AngII concentrations were not different between hepAGT +/+ and hepAGT -/- mice, which were also not affected by infection of AAVs encoding mouse AGT(L11V;Y12I) (Figure 3C). Renal AngII concentrations were higher in hepAGT +/+ mice and hepAGT -/- mice infected with mouse AGT(L11V;Y12I).AAV than in hepAGT -/- mice infected with null AAV (Figure 3D). Plasma total cholesterol concentrations were not different among the 3 groups (Figure II in the online-only Data Supplement). Population of mouse AGT(L11V;Y12I) in hepAGT -/- mice led to increased systolic blood pressure (Figure 3E) and atherosclerotic lesion size (Figure 3F), which were comparable to the two phenotypes in hepAGT +/+ mice infected with null AAV. A separate study demonstrated that female hepAGT -/- mice populated with mouse AGT(L11V;Y12I) had comparable blood pressure and atherosclerotic lesion size as female hepAGT +/+ mice injected with null AAV (Figure III in the online-only Data Supplement). We also compared hepAGT -/- female mice infected with wild type mouse AGT.AAV versus mouse AGT(L11V;Y12I). AAV. With same genome copies of AAVs, population of wild type AGT and mouse AGT(L11V,Y12I) had comparable elevations of plasma AGT concentrations, which were lower than hepAGT +/+ mice injected with null AAV (Figure III in the online-only Data Supplement). Despite the profound differences of plasma AGT concentrations, the 3 groups had comparable blood pressure and atherosclerotic lesion size.

**Figure 3.**
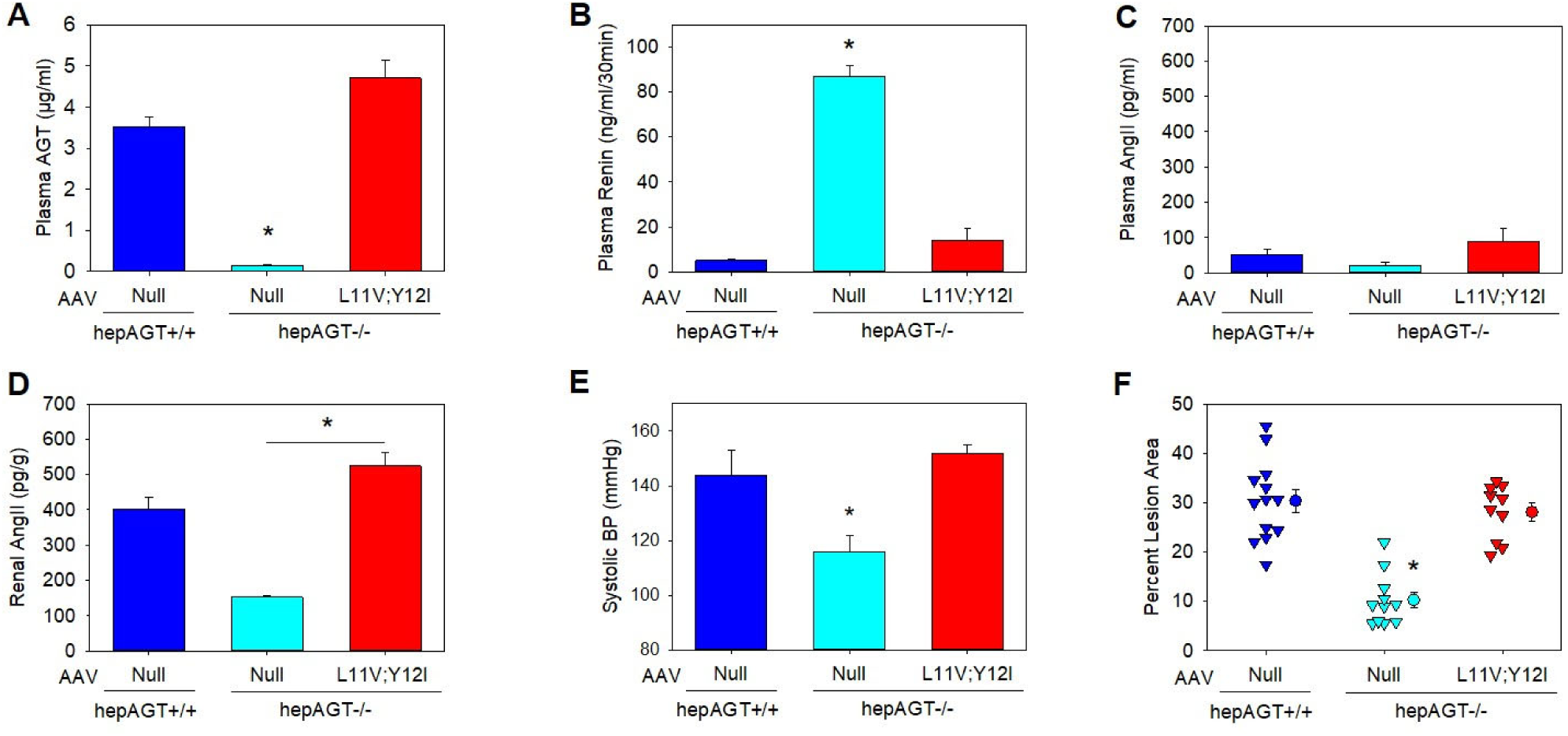
Mouse AGT(L11V;Y12I) restored Angll-mediated functions in male hepAGT -/- mice. **(A)** Plasma mouse AGT concentrations were measured by ELISA. *P<0.05 versus the other 2 groups by Kruskal–Wallis one way ANOVA on Ranks with Dunn’s method. **(B)** Plasma renin concentrations were measured by radioimmunoassay. *P<0.05 versus the other 2 groups by Kruskal–Wallis one way AISIOVA on Ranks with Dunn’s method. Angll concentrations in plasma **(C)** and kidney **(D)** were measured using LC-MS/MS. *P<0.05 by Kruskal–Wallis one way ANOVA on Ranks with Dunn’s method. **(E)** Systolic blood pressure was measured using a tail-cuff system. *P<0.05 versus the other 2 groups by one way ANOVA with Holm-Sidak method. **(F)** Atherosclerotic lesion area was quantified by en face method. *P<0.05 versus the other 2 groups by one way ANOVA with Holm-Sidak method. N = 10 - 13/group.

## DISCUSSION

Using an AAV approach for manipulating AGT in hepatocytes, this study reports two novel findings. First, human AGT does not affect AngII-mediated functions in mice lacking endogenous AGT in hepatocytes. Second, replacement of the amino-terminal amino acids at positions 11 and 12 of mouse AGT to mimic human AGT does not impair renin cleavage of AGT, resulting in normal renal AngII production and AngII-mediated functions.

Previous studies using human AGT transgenic mice have demonstrated that the presence of human AGT in mice does not affect systolic blood pressure.^15-17^ In human AGT transgenic mice, human AGT was expressed in mice with normal concentrations of endogenous mouse AGT and relatively low mouse renin concentrations, which does not rule out the possibility that lack of cleavage of human AGT is due to low mouse renin concentrations. Our mouse model has several advantages: (1). We used a mouse model having low endogenous mouse AGT concentrations but high mouse renin concentrations. The low plasma concentrations of mouse AGT were sufficient to enable normal growth and kidney development, but led to much lower blood pressure, compared to their wild type littermates.^4,9,10^ Therefore, this mouse model is relevant to physiological regulation of the renin-angiotensin system. More importantly, our mouse model can address the caveats raised above in previous studies. (2). Our mouse model was on an LDL receptor -/- background, so we were able to study two different AngII-induced physiologic and pathophysiologic functions, namely blood pressure and atherosclerosis. (3) The AAV vector encoding human AGT with a hepatocyte-specific promoter led to human AGT produced in hepatocytes, mimicking the normal processing of AGT production in liver and secretion to the circulation.^10,18^

In hepAGT -/- mice infected with AAV containing human AGT, plasma human AGT concentrations were above 10 μg/ml, which were 3 - 4-fold higher than plasma endogenous mouse AGT concentrations in wild type mice. Although mouse plasma renin concentrations were greatly elevated in hepAGT -/- mice compared with wild type mice due to removal of AngII-mediated negative feedback, the presence of human AGT did not correct plasma renin concentrations, elevate blood pressure, or contribute to the development of atherosclerosis. Therefore, expression of human AGT did not restore AngII-mediated functions in hepAGT -/- mice, which supports the notion that human AGT is not cleaved by mouse renin in vivo.^5-7,15^ These results confirm previous findings of species specificity of AGT and renin interaction,^5,6,19^ and extend these findings to the in vivo physiological and pathophysiological setting.

In contrast to population of human AGT in hepAGT-/- mice, population of mouse AGT with L11 and Y12 being substituted to the residues of human AGT reduced plasma renin concentrations and increased renal AngII concentrations to those of wild type mice. As anticipated with these findings, the responses of blood pressure and atherosclerotic lesion size were restored in hepAGT -/- mice infected with murine AGT containing these two residues of human AGT. These findings provide compelling evidence that these two amino acid substitutions between murine and human do not affect renin cleavage of AGT and the subsequent AngII-mediated functions in hepAGT -/- mice. Our results do not support a role for these two residues of human AGT as determinants of the species-specificity of the reaction between AGT and renin. In addition to amino acid sequences, many other conditions may affect AGT binding and cleavage by renin such as AGT protein tertiary structure. Mapping conserved sequences among species on AGT tertiary protein structure may facilitate the definition of residues that determine renin cleavage efficacy.^2,3^

Consistent with our previous report,^10^ plasma AngII concentrations were not different between hepAGT +/+ and -/- mice, irrespective of infection with AAV containing a null insert or mutated AGT. In contrast, population of mouse AGT with L11V;Y12I substitutions increased renal AngII concentrations to levels of hepAGT +/+ mice, findings consistent with changes of blood pressure and atherosclerosis in these mice. In this study, AngII concentrations in kidney were considerably higher than in plasma. AT1a receptor is abundant in kidney.^20^ Indeed, previous studies have reported that expression of AT1a receptor in kidney, but not in the vasculature, contributes to blood pressure regulation by AngII.^21,22^ Since renal AngII, but not plasma AngII, reflected AngII effects on blood pressure and atherosclerosis, these results suggest that local tissue AngII (e.g. renal) is important in these effects of the renin-angiotensin system.

We demonstrated same results in female mice, where mouse AGT(L11V;Y12I), but not human AGT, increased both blood pressure and atherosclerosis of hepAGT -/- mice to those of hepAGT +/+ mice. These results suggest that biologic sex is not a determinant of these findings.

In summary, the amino-terminal amino acids at positions 11 and 12 of human AGT that are adjacent to the renin cleavage site, do not determine renal AngII production and AngII-mediated functions in mice. Given the species-specificity of AGT cleavage by renin, we hypothesize that differences of either other amino acid sequences or the protein structure of AGT affect the renin cleavage. Future studies to identify species specificity of renin cleavage of AGT should align AGT protein sequences from different species and map the sequences to AGT tertiary structures.

## ACKNOWLEDGMENTS

We thank Victoria English for measuring plasma renin concentrations.

## SOURCES of FUNDING

The authors’ research work was supported by the National Heart, Lung, and Blood Institute of the National Institutes of Health under award numbers R01HL139748. The content in this manuscript is solely the responsibility of the authors and does not necessarily represent the official views of the National Institutes of Health.

## DISCLOSURES

None.

---

**Figure I.**
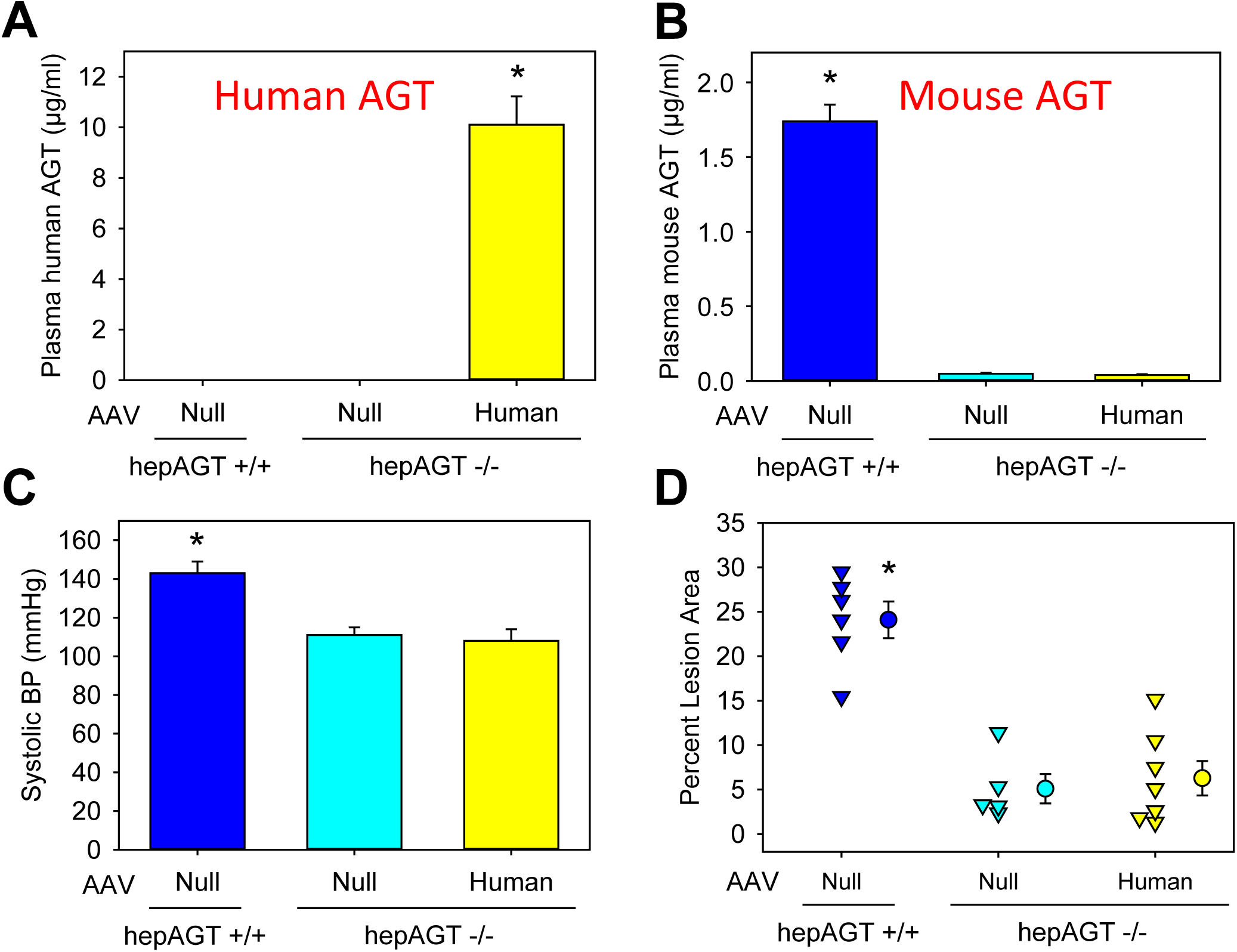
Human AGT did not affect AngII-mediated functions in female hepAGT-/- mice. Plasma human (**A**) and mouse (**B**) AGT concentrations were measured by ELISA. *P<0.05 versus the other groups by Kruskal–Wallis one way ANOVA on Ranks with Dunn’s method. (**C**) Systolic blood pressure was measured using a tail-cuff system. *P<0.05 versus the other groups by one way ANOVA with Holm-Sidak method. (**D**) Atherosclerotic lesion area was quantified by an en face method. *P<0.05 versus the other groups by one way ANOVA with Holm-Sidak method. N = 5 − 7/group.

**Figure II.**
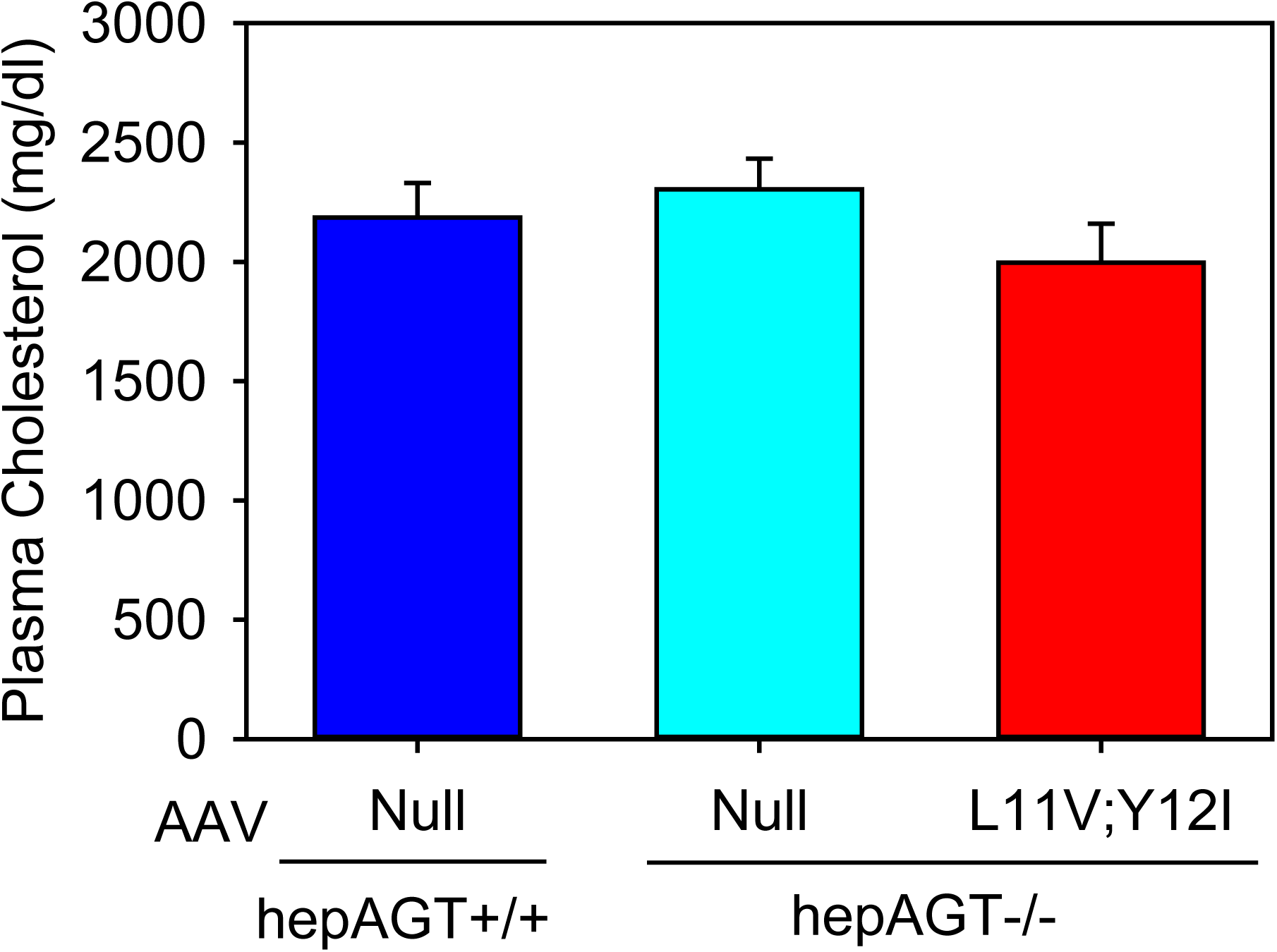
Mouse AGT(L11V;Y12I) did not affect plasma cholesterol concentrations in male mice. Male hepAGT-/- mice were injected intraperitoneally with null AAV or L11V;Y12I AGT AAV. hepAGT+/+ mice injected with null AAV were served as a control. Plasma total cholesterol concentrations were measured using an enzymatic kit and analyzed by one way ANOVA. N = 10 − 13/group.

**Figure III.**
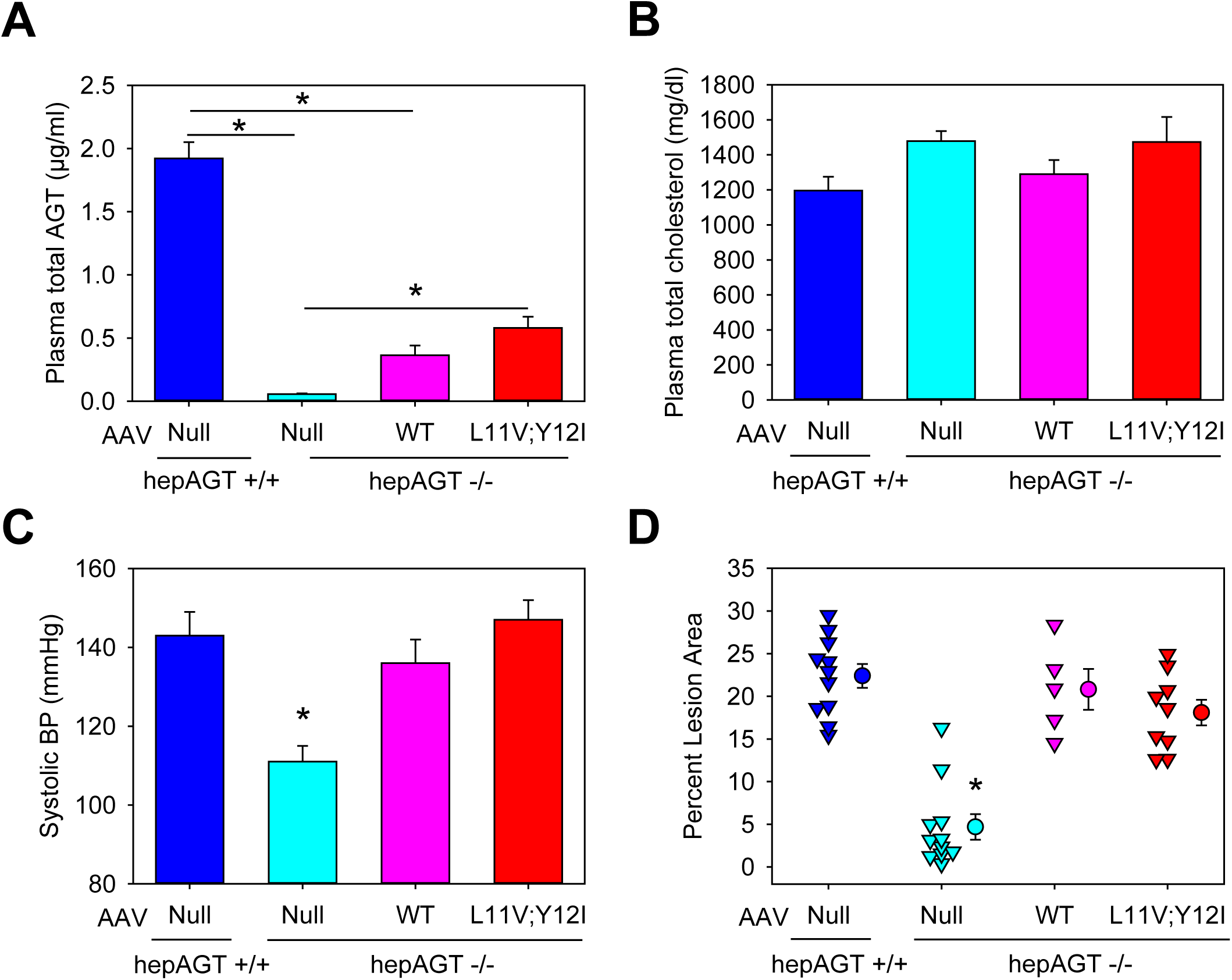
Mouse AGT(L11V;Y12I) had comparable effects to wild type mouse AGT on AngII-mediated functions in female mice. (**A**) Plasma mouse AGT concentrations were measured by ELISA. *P<0.05 by Kruskal–Wallis one way ANOVA on Ranks with Dunn’s method. (**B**) Plasma total cholesterol concentrations were measured using an enzymatic kit and analyzed by one way ANOVA. (**C**) Systolic blood pressure was measured using a tail-cuff system. *P<0.05 versus the other groups by one way ANOVA with Holm-Sidak method. (**D**) Atherosclerotic lesion area was quantified by an en face method. *P<0.05 versus the other groups by one way ANOVA with Holm-Sidak method. N = 5 − 11/group.

## DETAILED METHODS

### Animals

Development of hepatocyte-specific AGT deficient mice has been reported previously.^1-3^ AGT floxed (termed “Agt f/f”) x albumin-Cre^-/-^ (hepAGT +/+) and Agt f/f x albumin-Cre^+/-^ (hepAGT -/-) littermates were used for experiments described in this manuscript. Genotypes were determined by Cre PCR and confirmed by measuring plasma AGT concentrations. Both male and female mice in an LDL receptor -/- background were studied following the recent ATVB Council statement.^4^ All animal experiments reported in this manuscript were performed with the approval of the University of Kentucky Institutional Animal Care and Use Committee (IACUC protocol number 2006-0009 or 2018-2968).

### Production and injection of adeno-associated viral (AAV) vectors

AAV vectors (serotype 2/8) driven by a hepatocyte-specific thyroxine-binding globulin (TBG) promoter were produced by the Vector Core in the Gene Therapy Program at the University of Pennsylvania. Three AAV vectors were made: (1) a null insertion (null AAV; used as control), (2) encoding the human AGT (human AGT.AAV), and (3) encoding the mouse AGT with L11V;Y12I mutations (L11V;Y12I.AAV). The L11V;Y12I represented that the N-terminal leucine at position 11 and tyrosine at position 12 in mouse AGT were replaced by valine and isoleucine, respectively, to mimic the two amino acids at the same positions in human AGT.

All mice were fed a normal laboratory diet (Diet # 2918; Envigo), and were injected intraperitoneally with AAV vectors (3 × 10^10^ genome copies/mouse). hepAGT +/+ mice were injected with null AAV as a positive control at 8 - 12 weeks of age. Sex- and age-matched hepAGT-/- mice were randomized to receive AAVs containing a null insert (negative control), human AGT, or L11V;Y12I mouse AGT. Two weeks after injection, mice were fed a diet supplemented with saturated fat (milk fat 21% wt/wt) and cholesterol (0.2% wt/wt; termed “Western diet”; Diet # TD.88137; Envigo) for 12 weeks (Figure I in the online-only Data Supplement).

### Plasma profiles

Mouse blood were collected in the presence of EDTA (final concentration: 1.8 mg/ml) and a proteinase inhibitor cocktail (provided by Attoquant Diagnostics GmbH, Vienna, Austria) to prevent degradation of AGT and angiotensin peptides. During the experiment, blood was collected through retro-orbital bleeding at selected intervals. Cardiac bleeding via right ventricle was used to collect blood at termination.

Plasma AGT concentrations were measured using a mouse AGT ELISA kit (Code # 27413; IBL America) or a human AGT ELISA kit (Code # 27412; IBL America).

Plasma renin concentrations were measured using a radioimmunoassy method:^5^ Mouse plasma samples (8 μl) were incubated in an assay buffer (Na_2_HPO_4_ 0.1 M, EDTA 0.02 M, maleate buffer pH 6.5, phenylmethyl-sulfonyl fluoride 2 μl; total volume of 250 μl) with an excess of rat AGT at 37 °C for 30 minutes. The reaction was terminated by placing samples at 100 °C for 5 minutes. AngI generated in each sample was quantified by radioimmunoassay using a commercially available kit (Cat # 1553; DiaSorin).

Plasma total cholesterol concentrations were measured using an enzymatic commercial kit (Cat # 999-02601; Wako Chemicals USA).

### Systolic blood pressure measurements

Systolic blood pressure was measured on conscious mice using a non-invasive tail-cuff system (Coda 8, Kent Scientific Corporation) following our standard protocol.^6^ Data were collected and analyzed based on 20 measurements of each mouse every day for 3 consecutive days. Mean systolic blood pressure of each mouse from the 3-day measurements was used for data analysis.

### Quantification of atherosclerosis

Atherosclerosis was quantified on the aortic intima including the ascending region, aortic arch, and 3 mm of the descending region using an en face method following the recommendations of the recent AHA statement.^7^

### Angiotensin II (AngII) Measurements

Plasma and renal AngII concentrations were measured using liquid chromatography–mass spectrometry by Attoquant Diagnostics GmbH (Vienna, Austria). All samples were assayed in a blinded manner.

